# Anticipatory Biogenesis of Hepatic Fed MicroRNAs is Regulated by Metabolic and Circadian Inputs

**DOI:** 10.1101/2023.04.17.537209

**Authors:** U.S. Sandra, Shreyam Chowdhury, Ullas Kolthur-Seetharam

**Affiliations:** Department of Biological Sciences, Tata Institute of Fundamental Research, Mumbai, India; Tata Institute of Fundamental Research- Hyderabad (TIFR-H), Hyderabad, India; Lead Contact

**Keywords:** Molecular Anticipation, Fed-Fast cycles, Physiology, Starvation, Micro-RNA.

## Abstract

Starvation and refeeding are mostly unanticipated in the wild in terms of duration, frequency, and nutritional value of the refed state. Notwithstanding this, organisms mount efficient and reproducible responses to restore metabolic homeostasis. Hence, it is intuitive to invoke expectant molecular mechanisms that build anticipatory responses to enable physiological toggling during fed-fast cycles. In this regard, we report anticipatory biogenesis of oscillatory hepatic microRNAs, which were earlier shown to peak during a fed state to inhibit starvation-responsive genes. Results presented in this study clearly demonstrate that the levels of primary and precursor microRNA transcripts increase during a fasting state, in anticipation of a fed response. We delineate the importance of both metabolic inputs and circadian cues in orchestrating microRNA homeostasis in a physiological setting, using the most prominent hepatic fed-miRNAs as candidates. Besides illustrating the metabo-endocrine control, our findings provide a mechanistic basis for the overarching influence of starvation on anticipatory biogenesis. Importantly, by employing pharmacological agents that are widely used in the clinics, we point out the high potential of interventions to restore homeostasis of hepatic microRNAs, whose deregulated expression is otherwise well established to cause metabolic diseases.

## INTRODUCTION

Fed-fast cycles and the associated anabolic-catabolic transitions are vital for physiological homeostasis and inefficient toggling is known to cause metabolic diseases and accelerated aging (Ribarič, 2012; Suliga *et al*., 2015; Geisler *et al*., 2016; Petersen, Vatner and Shulman, 2017; Paoli *et al*., 2019). Evolutionarily, these transitions are coupled with circadian oscillations and dictated by light-dark and sleep/inactive-wake/active cycles feed (Gerhart-Hines and Lazar, 2015; Pickel and Sung, 2020a). Physiological changes that accompany fed-fast and circadian cycles are dependent upon gene expression cascades especially in central metabolic tissues such as the liver (Vollmers *et al*., no date; Jitrapakdee, 2012; Bideyan, Nagari and Tontonoz, 2021). Even though decades of work have illustrated the congruence of nutrient and circadian inputs for gene transcription, if/how they impact post-transcriptional regulation of metabolic gene programs is relatively less understood.

MicroRNA (miR)-dependent gene expression control is well established and particularly important for mediating dynamic changes in mRNA translation, storage and turnover (Wilczynska and Bushell, 2015; O’Brien *et al*., 2018; Dexheimer and Cochella, 2020). We and others have demonstrated the significant role of miR-mediated regulation of gene expression in orchestrating physiological responses including during fed-fast cycles (Hu *et al*., 2012; Rottiers and Näär, 2012; Maniyadath *et al*., 2019; LaPierre *et al*., 2022). In addition to specific miR signatures that are associated with fed and fasted states in the liver (Na *et al*., 2009; Vollmers *et al*., 2012; Ji *et al*., 2023), hepatic miRs have also been shown to impinge on distant organ systems (Sung, Kim and Jung, 2018; Liu *et al*., 2020; Ji *et al*., 2021). However, metabolic inputs/signals that are essential for miR-homeostasis, from biogenesis and processing to degradation and secretion, are largely unknown. This is likely the case since, as detailed later, activities of drosha and dicer have been shown to be regulated in a few other physiological contexts (Davis and Hata, 2009; Creugny, Fender and Pfeffer, 2018). Moreover, given that fed-fast cycles are inherently linked to circadian rhythm, there is a paucity of information vis-à-vis phenomenological and mechanistic underpinnings of hepatic miR biogenesis and processing.

Decades of work have demonstrated that organismal behaviour and physiology, which are associated with feeding and fasting involve anticipation, and are largely mediated by cephalic responses (Power and Schulkin, no date a; Zafra, Molina and Puerto, 2006; Smeets, Erkner and De Graaf, 2010). Cephalic responses are conditioned hormonal and metabolic adaptations, and perturbing such anticipatory responses have been shown to have detrimental effects (Ahré and Holst, 2001; Zafra, Molina and Puerto, 2006). For example, blocking the cephalic phase of insulin secretion causes dysregulated glucose homeostasis (Ahré and Holst, 2001). While the current understanding of fed-fast anticipation is dominated by cephalic mechanisms that operate at an organismal level, it is still unclear if anticipatory cellular-level mechanisms facilitate effective switching between anabolic and catabolic states. Further, whether such cellular/molecular anticipatory mechanisms are tuned by neuro-endocrine and metabolic signals remains unknown.

It is interesting to note that the hepatic fed microRNAs that we previously characterized as regulators of hepatic functions and organismal physiology exhibited anticipatory biogenesis (Maniyadath *et al*., 2019). Specifically, the dynamic post-transcriptional convergent and additive control exerted by the fed-miR network entailed the transcription of these microRNAs in a fasted/starvation state. Among these microRNAs, let-7i, miR-221, miR-222, and miR-204 have now emerged as key players in various liver pathological conditions including fibrosis and hepatocellular carcinoma (Chen *et al*., 2014; Luo *et al*., 2017; Song *et al*., 2017a; Di Martino *et al*., 2022; Kim *et al*., 2022). Therefore, identifying the upstream mechanisms that govern their biogenesis will not only provide fundamental insights into molecular anticipation but also possibly open up new avenues for therapeutic interventions.

It is intriguing to note that despite the high volume of information vis-à-vis miR-dependent regulation of cellular and organismal functions (Ebert and Sharp, 2012; Dexheimer and Cochella, 2020), including metabolism and physiology (Rottiers and Näär, 2012; Vienberg *et al*., 2017), our understanding of mechanisms that dictate miR homeostasis is still poor. Seminal studies have illustrated that Ras/MAPK and TGF-/BMP signaling pathways play a major role as regulators of microRNA biogenesis (Hata and Davis, 2009; Kent *et al*., 2010; Saj and Lai, 2011). Even though studies have also unravelled specific transcription factors that are necessary for miR expression (Davis and Hata, 2009; Saj and Lai, 2011; Creugny, Fender and Pfeffer, 2018), little is known about differential and/or layered control of miR processing that together determine biogenesis. For example, expression and activities of drosha and dicer have been shown to be regulated in response to several stresses (Davis and Hata, 2009; Beezhold, Castranova and Chen, 2010; Creugny, Fender and Pfeffer, 2018). Albeit, these reports in conjunction with emerging evidence on RNA methylation, portend layered regulation of miR homeostasis, the physiological significance of such regulation particularly in the context of hepatic fed-fast transitions is unknown as yet. Moreover, although hepatocyte nuclear factor 4α (HNF4α) and peroxisome proliferator-activated receptor α (PPARα) have been shown to transcribe miR-genes in hepatocytes (Li *et al*., 2011; Cheng *et al*., 2017), it will be important to investigate if these are orchestrated by endocrine and metabolic signals to create molecular anticipation vis-à-vis fed-miR biogenesis.

In the present study, we clearly illustrate the anticipatory expression of hepatic fed microRNAs during a starvation state. We provide phenomenological and mechanistic underpinnings to delineate the contributions of fed-fast and circadian inputs in eliciting this anticipatory expression, besides providing the complex regulation of primary, precursor and mature microRNA transcripts in the liver.

## MATERIALS AND METHODS

### Animal Experiments

Animal experiments were performed using C57BL6NCrl mice that were reared under standard animal house conditions (12-hour light-dark cycle and fed a regular chow diet). 2-3 months old male mice were used for all diet and circadian perturbations, unless specified otherwise. For aged cohorts 20–22 months old male mice were used. considered as young and aged cohorts, respectively. The procedures and the project were approved and were in accordance with the Tata Institute of Fundamental Research (TIFR) Institutional Animal Ethics Committee (IAEC) (CPCSEA-56/1999).

Fed-starved-refed paradigms: (a) Fed mice – *ad libitum* fed mice sacrificed at ZT1 (8.00 am) (b) Starved mice – *ad libitum* fed mice starved for 24 hours and sacrificed at ZT1 (8.00 am) (c) Refed mice – post 24 hours of starvation mice were fed *ad libitu*m for 6 hours and sacrificed at ZT7 (2.00 pm). Both young (2-3 months old) and aged (20-22 months old) male mice were used for these experiments as indicated.

Light-fed and dark-fed paradigms: mice were entrained for 14 days for time restricted feeding with exclusive access to food either during the light phase (ZT24 – ZT12) or the dark phase (ZT12 – ZT24), as indicated.

Light-dark *ad libitum* (LD-AL) and starvation (LD-S) paradigms: mice were subjected to starvation by removing the feed at ZT0 (7 am), designated as LD-S. Mice that were *ad libitum* fed served as controls, designated at LD-AL. Both LD-S and LD-AL mice were euthanized at same ZT time points at 4-hour interval (starting from ZT4).

Dark-dark paradigms: mice housed under constant darkness for 24 hours were subjected to different durations of starvation by removing feed at ZT0, designated as DD-S. DD-S mice were compared with those that had *ad libitum* access to feed during the continuous dark phase, designated as DD-AL. In both cases animals were euthanized at 4-hour intervals (as indicated) and all mice handling and tissue harvesting were done under dim red light.

High-fat diet (HFD) and sucrose supplementation paradigms: (a) mice were provided *ad libitum* access to normal chow or 35% High Fat diet for 12 weeks and (b) normal chow diet ad libitum fed mice were given access to only water or 10% sucrose in water for 12 weeks. For both the paradigms, mice were starved for 12 hours starting at ZT13 (8 pm) and sacrificed at ZT1 (8 am).

### RER measurements

For RER measurement, O_2_ consumption, and CO_2_ production were measured, using indirect calorimetry in an Oxymax Comprehensive Lab Animal Monitoring System CLAMS open circuit system (Columbus Instruments, Columbus, OH, USA). Animals were housed in individual metabolic cages with a constant airflow of 0.5 L/min and under standard conditions with *ad libitum* access to food and water. Readings from each cage were measured every 20 min for 48 hours (12h light/12h dark cycle). The respiratory exchange ratio (RER) was derived from the measured volume of oxygen consumed (VO_2_) and carbon dioxide produced (VCO_2_) that were normalized to body mass (mL/kg/h).

### Primary Hepatocyte Isolation

10–12 weeks old male mice were used for primary hepatocyte isolation. The liver was perfused with Buffer A (HBSS, 1 mg/ml D-glucose, 25 mM HEPES, and 0.5 mM EGTA) and digesting medium (low glucose DMEM (LG, 5 mM glucose), antibiotic antimycotic solution (AA), 15 mM HEPES, and 19 mg collagenase type IV) in a sequential manner. The perfused livers were then harvested and minced in digestion media. It was further passed through a 70-micron pore-size filter and centrifuged at 50 g for 5 minutes at 4°C to obtain the hepatocyte pellet.

### Primary Hepatocyte Culture

The hepatocyte pellet was then washed three times with high glucose DMEM media (HG, 25 mM glucose), followed by plating into cell culture plates (coated with collagen) in HG media containing 10 % fetal bovine serum (FBS). After six hours of plating, the culture media was replaced with serum-free HG media and left overnight. For assessing the autonomous cue-dependent miR biogenesis, hepatocytes were treated with high glucose-HG (25 mM), low glucose-LG (5 mM) or glucose-free media-NG for 12 hr. 2 mM metformin treatment was done in high glucose media for 12 hr. For assessing the endocrine-dependent miR biogenesis cells were treated with 30 nM glucagon in LG media for 6 hr, and 10 μM forskolin in LG media for 12 hr. For PPARα dependent regulation of miR expression, cells were incubated in HG media containing 50 μM WY-14643 for 10 and 24 hrs and 100 μM bezafibrate for 24 hrs. For anabolic cue dependent regulation of mature miRs cells were treated with HG media containing 100nM of insulin for 1.5 hrs. Following the treatments cells were collected in TriZol and proceeded for RNA isolation.

### RNA isolation, cDNA synthesis and quantitative PCR

RNA isolation, cDNA synthesis and qPCR were carried out as per manufacturer’s instructions. Briefly, total RNA was isolated from liver tissue/hepatocytes using TriZol reagent and 5 – 8 µg RNA was used to prepare cDNA using random hexamers and SuperScript-IV RT kit. Quantitative PCR (qPCR) was performed using KAPA SYBR® FAST Universal 2X qPCR Master Mix on Roche Light Cycler 480 II and LC 96 instruments. For mature microRNA profiling, the cDNA was prepared using The QuantiMir™ RT Kit, and qPCR were performed as per manufacturer’s instructions. For normalization, levels of actin mRNA/18S rRNA were used.

### Quantification and statistical analysis

Data are expressed as means ± standard error of means (SEM). Statistical analyses were done using Graph Pad Prism (version 8.0). Student’s t-test and ANOVA were used to determine statistical significance. A value of p ≤ 0.05 was considered statistically significant. *p ≤ 0.05; **p ≤ 0.01; ***p ≤ 0.001.

## RESULTS

### Diurnal oscillation in hepatic miR biogenesis

As reported earlier (Maniyadath *et al*., 2019), we found oscillatory expression of mature miRs in *ad libitum* fed, 24 hours starved and refed mice livers, as indicated (Fig 1A). Next, to address the diurnal rhythmicity of hepatic fed-microRNAs let-7i, miR-221, miR-222, and miR-204, we harvested livers from *ad libitum* fed mice sacrificed at 4-hour intervals, over the course of a 24-hour light-dark cycle. We found that these microRNAs oscillate in a diurnal fashion (Fig 1B) and correlated well with the circadian-dependent expression of *Cry1* and *Per2* (Fig S1A). Notably, their expression progressively decreased in the inactive light phase (ZT4- ZT12) and increased in the active dark phase (ZT16- ZT24) (Fig 1B). These time points corresponded with starved and fed states, as indicated by the Respiratory Exchange Ratio (RER) under *ad libitum* conditions (Fig S1B). The changes observed here were consistent with our earlier work (Maniyadath *et al*., 2019) and inversely corelated with the abundance of target starvation-dependent transcripts such as *Sirt1*, *Pgc1α*, *Lcad*, *Mcad*, and *Tfam* (Fig S1C). This clearly suggested that hepatic fed microRNAs follow dynamic oscillatory changes under *ad libitum* fed conditions.

**Figure 1:**
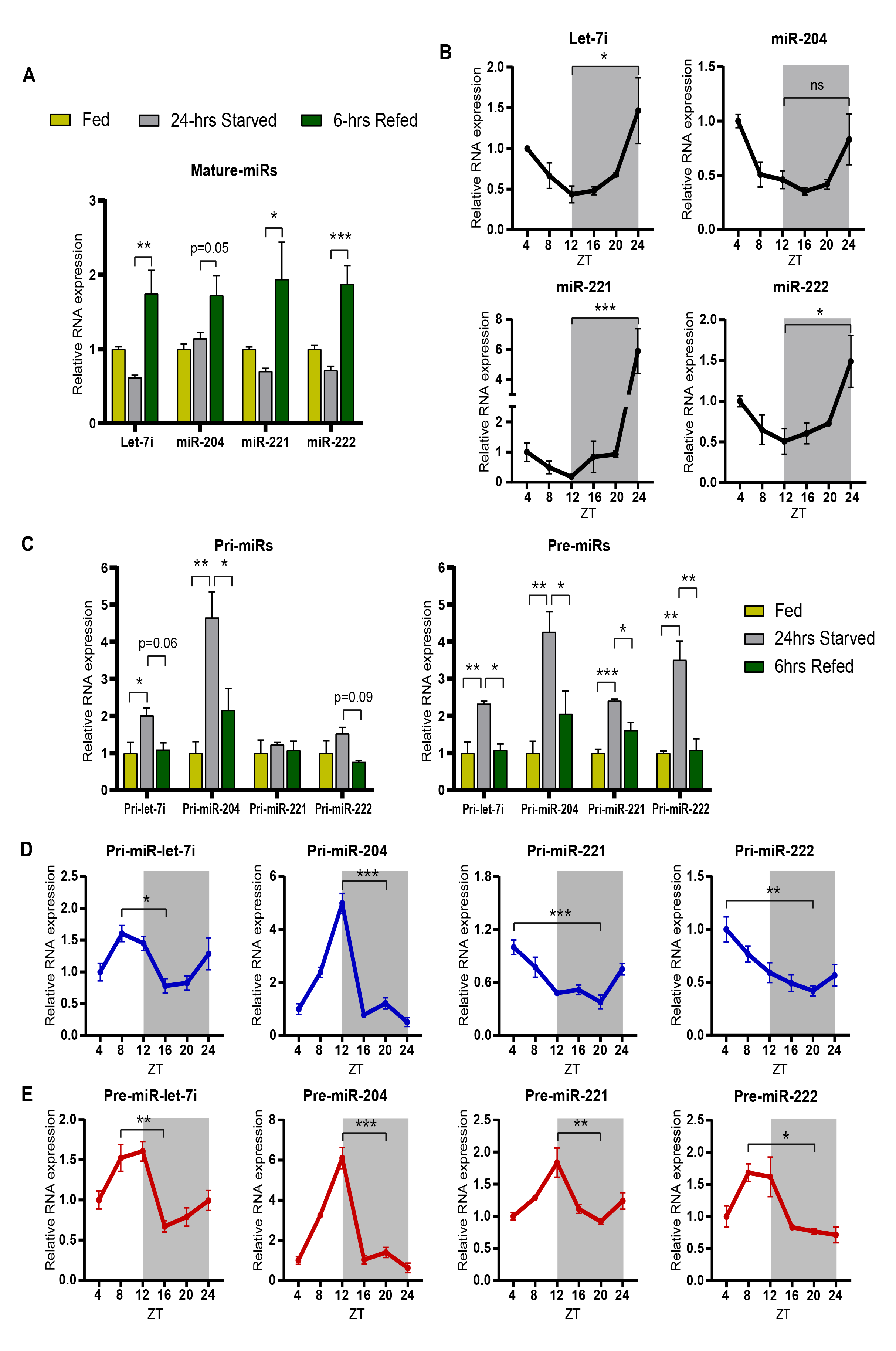
Diurnal oscillation of anticipatory microRNA-biogenesis in mouse liver. **(A)** Relative expression of mature miRNAs (miRs) - let-7i, miR-221, miR-222 and miR-204 in *ad libitum* fed, 24 hrs starved and 6 hrs refed mice. MiR transcript levels were normalized to 18S rRNA level in each case and plotted as fold change with respect to *ad libitum* fed condition (N=2, n=3). **(B)** Relative expression of mature miRs from liver of *ad libitum* fed mice at the indicated ZT (Zeitgeber time). MiR transcript levels were normalized to 18S rRNA level and plotted as fold change with respect to ZT4 (N=2, n=3). **(C)** Relative expression of primary miRNAs (pri-miRs) and precursor miRNAs (pre-miRs) in *ad libitum* fed, 24 hrs starved and 6 hrs refed mice. Pri-and pre-miR transcript levels were normalized to *Actin* mRNA level and plotted as fold change with respect to *ad libitum* fed (N=2, n=3). **(D-E)** Relative expression of **(D)** Pri-miR and **(E)** Pre-miR transcripts from liver of *ad libitum* fed mice at the indicated ZT. Pri-and pre-miR transcript levels were normalized to *Actin* mRNA level and plotted as fold change with respect to ZT4 (N=2, n=3). Data are represented as mean ± SEM. Statistical significance was calculated using one-way ANOVA with Tukey’s test for multiple comparisons between groups (*, p < 0.05; **, p < 0.01; ***, p < 0.001).

We had earlier demonstrated miR-dependent post-transcriptional inhibition of starvation genes upon refeeding and that this dynamic mode of control is possibly facilitated by anticipatory biogenesis of miRs during a fasted state (Maniyadath *et al*., 2019). Thus, we wanted to assess miR-biogenesis in *ad libitum* fed, 24 hrs starved, and 6 hrs refed mice. As expected, we observed an increase in primary and precursor transcripts of these microRNAs in 24 hr starvation, which was reduced or brought down to basal level upon refeeding (Fig 1C). This prompted us to investigate the oscillation of the primary and precursor transcripts across ZT time points in *ad libitum* fed mice.

Interestingly, both pri-and pre-transcripts of fed microRNAs were elevated in the starvation/inactive phase in a miR-specific manner (Fig 1D and E). For example, levels of pri-let-7i and pri-miR-204 exhibited a progressive increase from ZT4 onwards and peaked around ZT8-ZT12 (Fig 1D). While pri-miR-221 and pri-miR-222 also showed elevated levels in the light/starvation phase, their maximal expression was seen earlier at ZT4 (Fig 1D). Regardless of such differences in pri-miRs, pre-miRs showed similar patterns of induction during the light/starvation phase with a peak around ZT8-ZT12 (Fig 1E). Together, these results demonstrated that the pri-, pre-, and mature-transcripts of microRNAs let-7i, miR-221, miR- 222, and miR-204 showed diurnal rhythmicity. Further, they indicated a possible association between circadian and fed-fast cues in regulating miR homeostasis i.e., in addition to validating the anticipatory nature of miR biogenesis

### Temporally restricted feeding paradigms rewire miR biogenesis in the liver

It is well established that circadian oscillations and fed-fast cycles are intrinsically coupled (Pickel and Sung, 2020b). Specifically in the liver, nutrient/metabolic inputs have been shown to tune circadian dependent molecular rhythmicity and thus exert control over the peripheral clock (Vollmers *et al*., no date; Lamia, Storch and Weitz, 2008; Greenwell *et al*., 2019). Given this and based on the results presented above, we wanted to delineate between fed-fast and circadian dependence of anticipatory pri-miR and pre-miR expression. Towards this, we first employed a time-restricted feeding paradigm where food was made available either in the dark phase (ZT12- ZT24) (dark-fed) or in the light phase (ZT0-ZT12) (light-fed) (Fig 2A). Assaying for RER and *Cry1* expression, following the 14-day entrainment period, clearly showed the switch in whole body energetics and inverted circadian rhythm in the liver (Fig 2B and C), which was also correlated with expression of genes involved in starvation and hepatic metabolism (Fig 2D)

**Figure 2:**
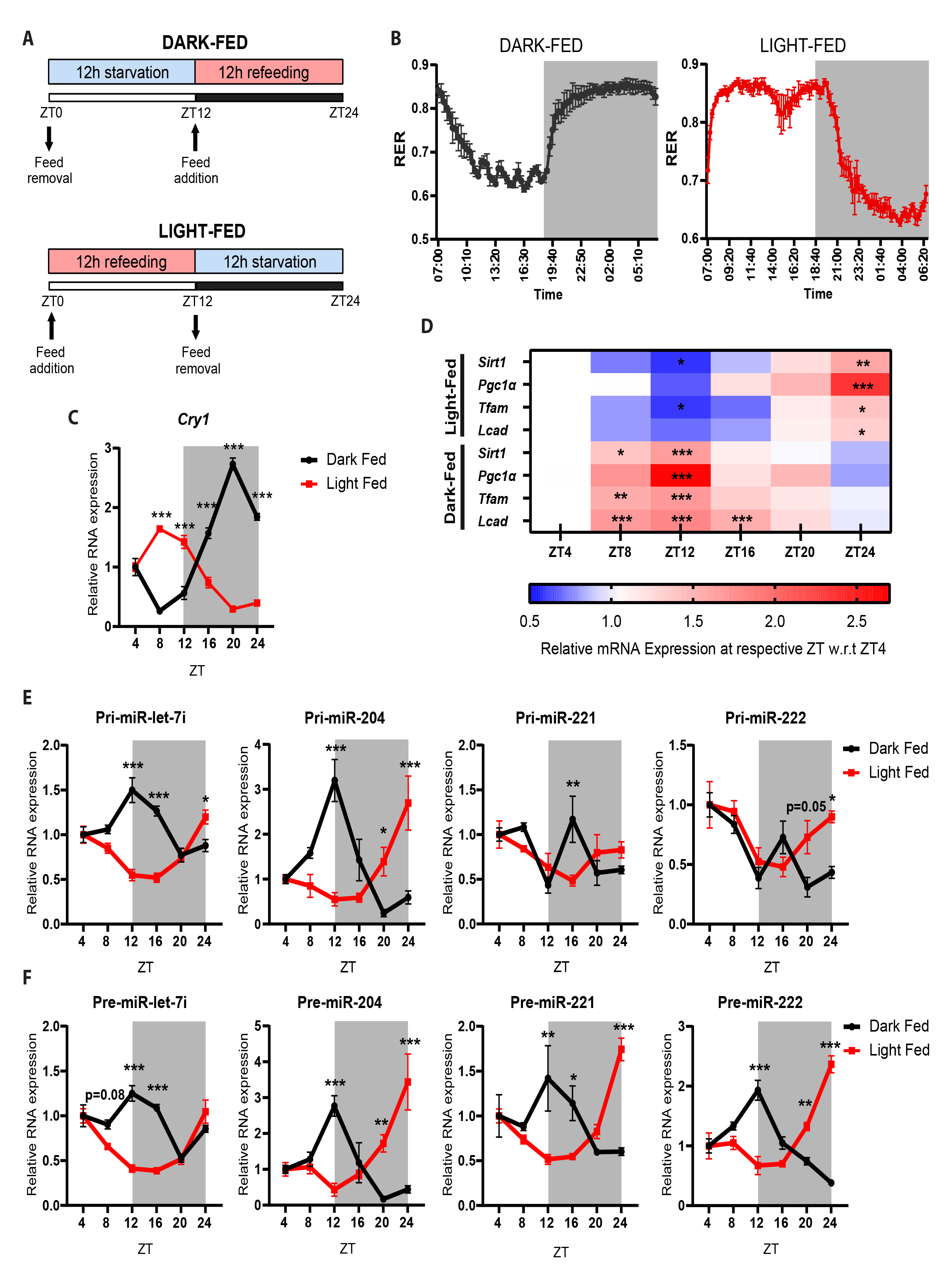
Temporal feeding paradigms reprogram the oscillatory miRNA-biogenesis in liver. **(A)** Schematic outline of the feeding paradigm employed. Dark-fed and light-fed mice were allowed access to food from ZT12-ZT24 and ZT0-ZT12 respectively. **(B)** Respiratory exchange ratio of dark-fed and light-fed mice over a 24-hr light-dark cycle (n=3). **(C)** Relative expression of *Cry1* mRNA from the liver of dark-fed and light-fed mice at the indicated ZT. *Cry1* transcript level was normalized to *Actin* mRNA level and plotted as fold change with respect to ZT4 for both dark-fed and light-fed mice (N=2, n=3). **(D)** Heatmap depicting relative expression of the starvation-responsive gene mRNAs at the indicated ZT. mRNA levels of the respective genes were normalized to *Actin* mRNA level and plotted as fold change with respect to ZT4 for both dark-fed and light-fed mice (N=2, n=3). **(E-F)** Relative expression of **(E)** Pri-miRs and **(F)** Pre-miRs from liver of dark-fed and light-fed mice at the indicated ZT. Pri-and pre-miR transcript levels were normalized *Actin* mRNA level and plotted as fold change with respect to ZT4 for both dark-fed and light-fed mice (N=2, n=3). Data are represented as mean ± SEM. For **C**, **E**, and **F**, statistical significance between dark-fed and light-fed group at indicated ZT was calculated using multiple t-tests with Holm-Sidak correction. For **D**, statistical significance was calculated using one-way ANOVA with Tukey’s test for multiple comparison at indicated ZT with respect to ZT4 (*, p < 0.05; **, p < 0.01; ***, p < 0.001).

Not surprisingly, the expression pattern of pri-, pre-, and mature-microRNA (Fig 2E-2F and Fig S2A) transcripts in the dark-fed group was similar to that of *ad libitum* fed mice and agreed well with fed-fast dependence. The only notable difference that we observed between *ad libitum* and dark-fed groups was for pri-miR-221 and pri-miR-222, which showed an additional peak at ZT16 that was absent in the *ad libitum* fed mice (Fig 2E). Importantly, while pri-let-7i and pri-miR-204 showed complete inversion in expression it was subdued for pri-miR-221 and pri-miR-222 when mice were fed during the light phase (Fig 2E). Irrespective of this it was interesting to observe that precursor transcripts of all the miRs assayed demonstrated retroverted expression (Fig 2F). Moreover, the maximal abundance of pre-miRs was seen at ZT12 and ZT24 in a paradigm-specific manner (Fig 2F), interestingly at time points that correlate with peak starvation and transition to dark or light phases, respectively.

Consistent with our hypothesis, we indeed found the levels of mature miRs were inversed in light-fed and dark-fed cohorts, with peak expression in fed states (Fig S2A) and as indicated above, anti-correlated with the target mRNAs. Although, we had earlier identified them as hepatic fed-miRs, we exploited the paradigms described here to investigate if fed endocrine inputs, in the form of insulin, were necessary to regulate the levels of mature miRs. Treating primary hepatocytes with insulin, as detailed in the methods section, led to an increase in mature miR-221 and miR-222 (Fig S2B). In addition to substantiating our earlier findings, these results clearly indicated the role of starvation and fed inputs in regulating the hepatic abundance of pri-, pre-and mature-microRNAs.

### Light independent regulation of let-7i/miR-204 but not miR-221/-222

Since both light-dark and fed-fast cycles are zeitgebers(Pickel and Sung, 2020b), we next wanted to check if miR biogenesis was independent of circadian/light-dark inputs. Towards this, we employed paradigms that perturbed both light-dark and fed-fast cycles (as detailed in the methods) and henceforth referred to as AL-LD (*ad libitum* light-dark paradigm) and S-LD (starvation light-dark paradigm) (Fig 3A). Specifically, to assess if anticipatory biogenesis was responsive to the duration/extent of starvation, we removed the feed at ZT0 and harvested livers from mice sacrificed at 4h intervals, as indicated (Fig 3A). While we found that the expression of clock and starvation-responsive metabolic genes were consistent with earlier reports (Fig S3A and B) (Vollmers *et al*., no date; Kinouchi *et al*., 2018), the primary and precursor transcripts of let-7i and miR-204 had lost oscillatory expression in response to continuous starvation, unlike in the *ad libitum* fed mice (Fig 3B and C). Notably, the decrease associated with the transition from the light to dark phase was absent (Fig 3B and C) and clearly hinted at transcriptional induction of let-7i and mir-204 independent of circadian inputs. In contrast, the response of pri-/pre-221/-222 was distinct and indicated an interplay between both fed-fast and circadian cues in regulating their homeostasis (Fig 3B and C). Albeit intriguing, the pattens displayed by pre-miR-221 and pre-miR-222 were disparate despite having common primary transcript, and corresponded well with studies that show vastly different levels of miR-221 and miR-222 (Wurz *et al*., 2010).

**Figure 3:**
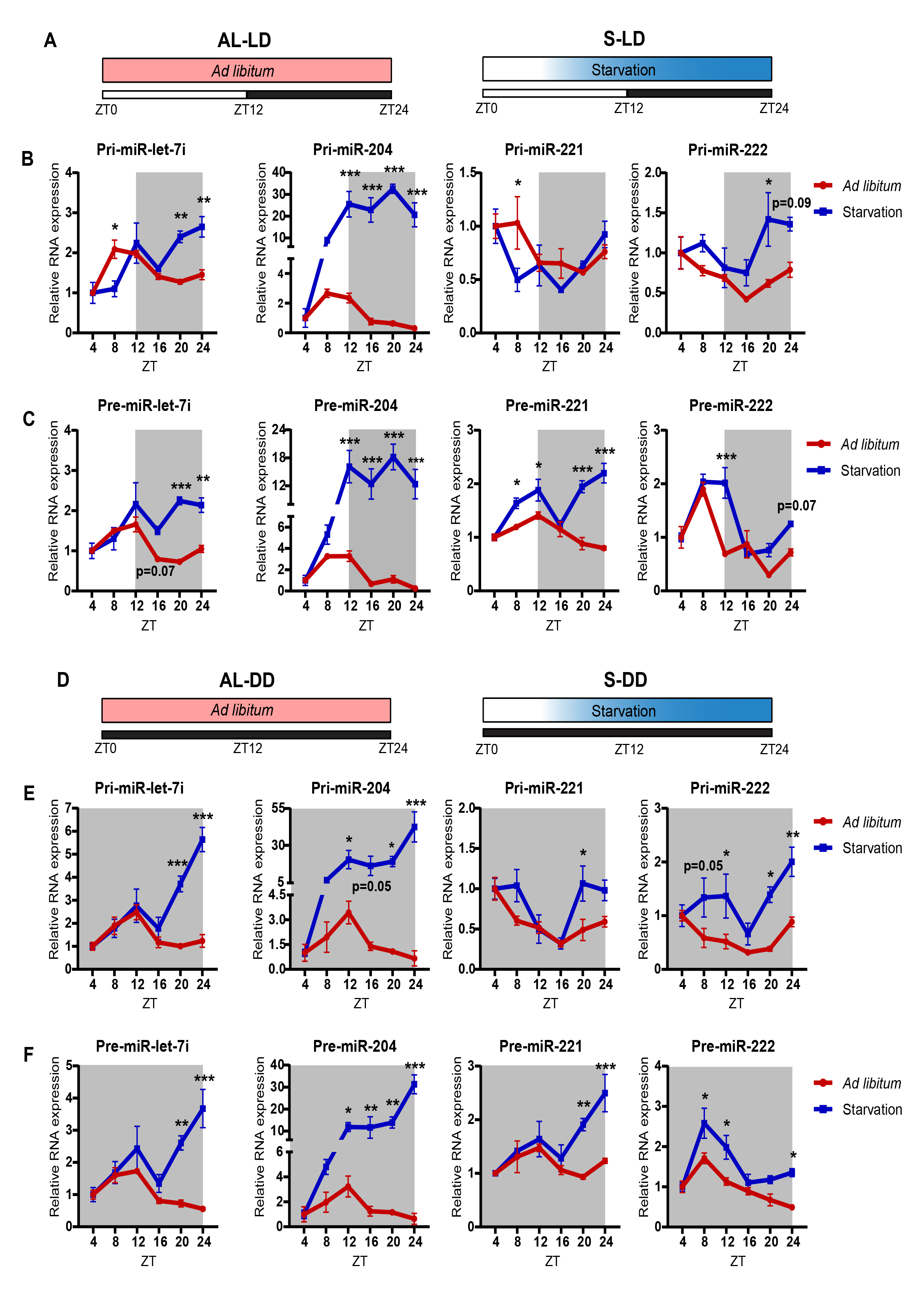
Hepatic miRNA biogenesis is responsive to progressive starvation and is independent of light inputs. **(A)** Schematic outline of the feeding paradigms employed. Mice were kept under a normal 12- hr light-dark cycle and either provided ad libitum access to feed (AL-LD) or starved from ZT0 (S-LD) for different durations (4, 8, 12, 16, 20 and 24 hrs) respectively. **(B-C)** Relative expression of **(B)** Pri-miRs and **(C)** Pre-miRs from the liver of AL-LD and S-LD mice, at the indicated ZT. Pri-and pre-miR transcript levels were normalized to *Actin* mRNA level and plotted as fold change with respect to ZT4 (N=2, n=3). **(D)** Schematic outline of the feeding paradigms employed. Mice were kept under constant darkness for 24 hrs and either provided ad libitum access to feed (AL-DD) or starved from ZT0 (S-DD) for different durations (4, 8, 12, 16, 20 and 24 hrs). **(E-F)** Relative expression of **(E)** Pri-miRs and **(F)** Pre-miRs from the liver of AL-DD and S-DD mice, at the indicated ZT. Pri-and pre-miR transcript levels were normalized to *Actin* mRNA level and plotted as fold change with respect to ZT4 (N=2, n=3). Data information: AL-LD - *ad libitum* light-dark, S-LD - starvation light-dark, AL-DD - *ad libitum* dark-dark, S-DD – starvation dark-dark, data are represented as mean ± SEM. Statistical significance between *ad libitum* fed and progressively starved mice at indicated ZT was calculated using multiple t-tests with Holm-Sidak correction (*, p < 0.05; **, p < 0.01; ***, p < 0.001).

To affirm these results and to provide conclusive evidence vis-à-vis dependence on either or both of the zeitgebers, mice housed in continuous dark cycle were subjected to starvation or fed *ad libitum* (S-DD and AL-DD, respectively), as indicated (Fig 3D). Interestingly, we found that the changes in expressions of the miRs assayed in AL-DD mice (Fig 3E and F) were nearly indistinguishable from AL-LD mice (Fig 3B and C). For example, primary and precursor transcripts of let-7i and miR-204 showed peak expression between ZT8 and ZT12, and a fall subsequently (Fig 3E and F). More importantly, levels of these pri-and pre-miRs in S-DD (Fig 3E and F) mice mirrored the pattern that was observed in S-LD (Fig 3B and C) mice. Together, these results clearly indicated that especially for let-7i and miR-204 starvation cues but not light dependent circadian inputs regulated their expression.

### MiR-biogenesis is perturbed in aged and over-nutrition mice liver

Others and we have earlier demonstrated the importance of transcriptional and post-transcriptional mechanisms in maintaining physiological homeostasis and causal effects of aberrant fed-fast cycles in aging and metabolic deficits (Ribarič, 2012; Suliga *et al*., 2015; Maniyadath *et al*., 2019; Bideyan, Nagari and Tontonoz, 2021; -S. Designed Research; T and -S. Performed Research; T, 2023). In this regard, we were tempted to profile pri-and pre-miRs in separate paradigms of over-nutrition and aging. It is interesting to note that we observed a dramatic reduction in the starvation dependent upregulation in miR biogenesis in aged mice when compared to the young mice (Fig 4A and B). Specifically, while starvation dependent induction of primary transcripts of let-7i and miR-204 was significantly dampened (Fig 4A and B), refed mediated reduction in pri-miR-221/-222 was lost in aged mice (Fig 4A). Importantly, fed-fast-refed oscillation in pre-miRs was conspicuously subdued in livers isolated from aged mice (Fig 4A and B).

**Figure 4:**
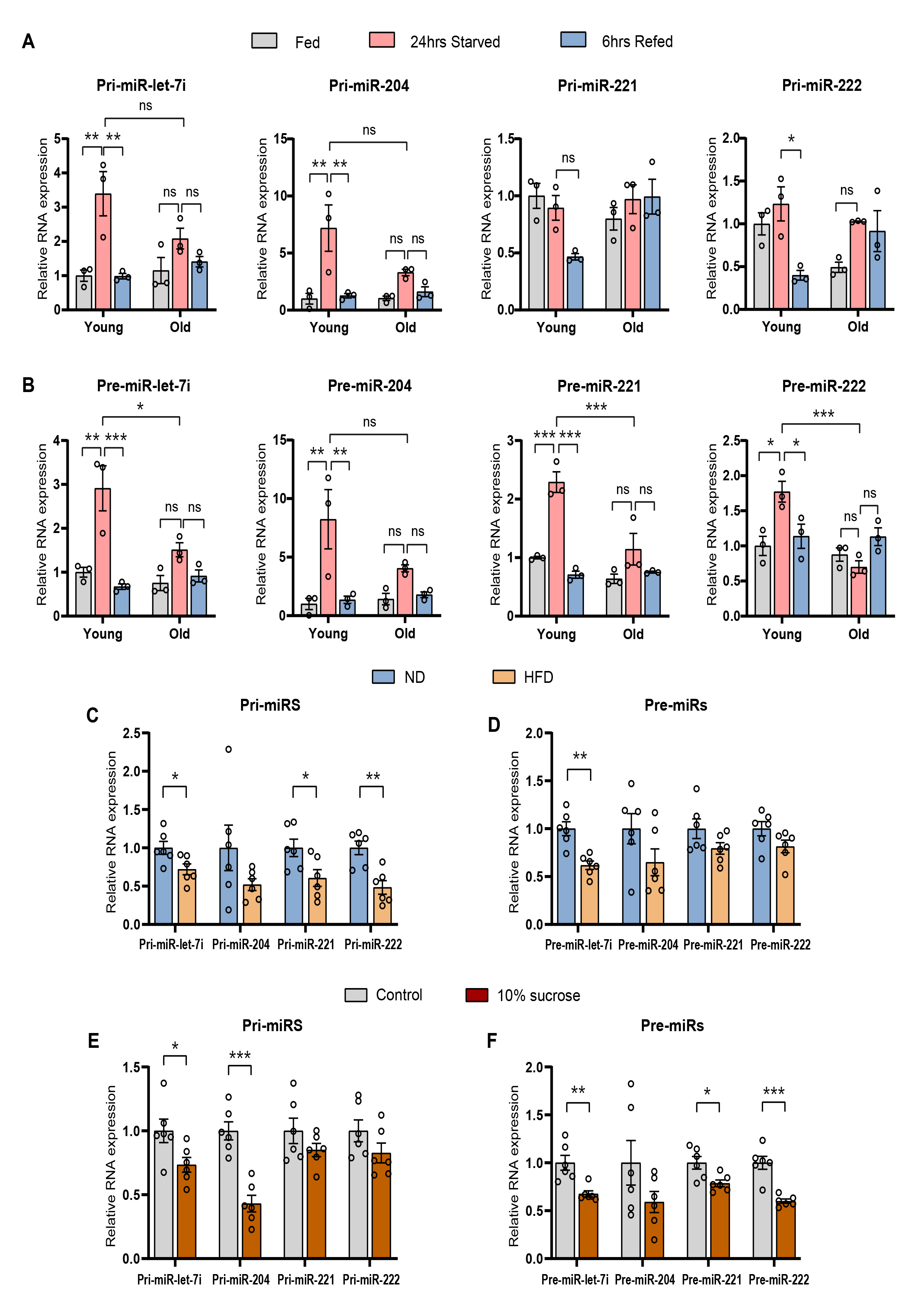
miRNA biogenesis is perturbed in aged and over-nutrition mice liver. **(A-B)** Relative expression of **(A)** Pri-miR and **(B)** Pre-miR in *ad libitum* fed, 24 hrs starved and 6 hrs refed young (2-4 months old) and old (20-22 months old) mice livers. Pri-and pre-miR transcript levels were normalized to *Actin* mRNA level and plotted as fold change with respect to *ad libitum* fed (N=2, n=3). **(C-D)** Relative expression of pri-and pre-miRs in 12 hrs starved mice liver subjected to 12 weeks of **(C)** 35 % HFD and **(D)** 10 % sucrose water (with regular chow). Pri-and pre-miR transcript levels were normalized to *Actin* mRNA level and plotted as fold change with respect to **(C)** Control diet and **(D)** Normal water (with regular chow) (N=2, n=3). Data are represented as mean ± SEM. For **A-B**, statistical significance between groups were calculated using two-way ANOVA with Tukey’s test for multiple comparisons. For **C-D,** statistical significance was calculated using a student t-test (*, p < 0.05; **, p < 0.01; ***, p < 0.001).

To study the effect of over-nutrition, we assayed for pri-/pre-miRs in livers harvested from mice fed with high-fat-diet (Fig 4C and D) and 10% sucrose in water (along with normal chow diet) (Fig 4E and F). We found diminished starvation response, in pri-/pre-miRs, in both these models when compared to chow-diet fed mice (Fig 4C, D, E and F). Overall, these results suggest that the anticipatory upregulation of miR biogenesis is lost/dampened under contexts, which are associated with perturbed physiological homeostasis.

### Starvation signals regulate miR-biogenesis via PPAR*α* in hepatocytes

Having demonstrated starvation dependence, we wanted to further investigate the molecular factors that mediate the upregulation of pri-/pre-miRs in hepatocytes. The hepatic starvation response is collectively governed by autonomous metabolic signalling/sensing and non-autonomous inputs predominantly from glucagon. In this regard, we assayed for primary and precursor miR transcripts in primary hepatocytes grown in medium containing differential glucose concentrations viz. 0mM (no glucose – NG), 5mM (low glucose – LG), and 25mM (high glucose – HG), to mimic fasting/fed states (Fig S4A).

It was interesting to find that while pri-/pre-miR-204 showed a robust upregulation in LG and NG, there was a mild induction in pre-miR-222, which indicated a partial glucose-deprivation dependent control (Fig S4A). Further, treating hepatocytes with glucagon, to evaluate the dependence on starvation endocrine inputs, resulted in elevated levels of pri-/pre-miRs (Fig 5A). To further substantiate these, we employed forskolin, a known activator of adenylate cyclase, which is downstream of the glucagon receptor, and observed a consistent induction that was akin to the response seen during starvation (Fig 5B). In order to pharmacologically mimic a starvation state that may also indicate the feasibility of therapeutic intervention, we treated primary hepatocytes with metformin, which is an FDA approved clinically administered drug to induce catabolic state. Interestingly, metformin treatment led to an increase in miR-biogenesis (Fig 5C).

**Figure 5:**
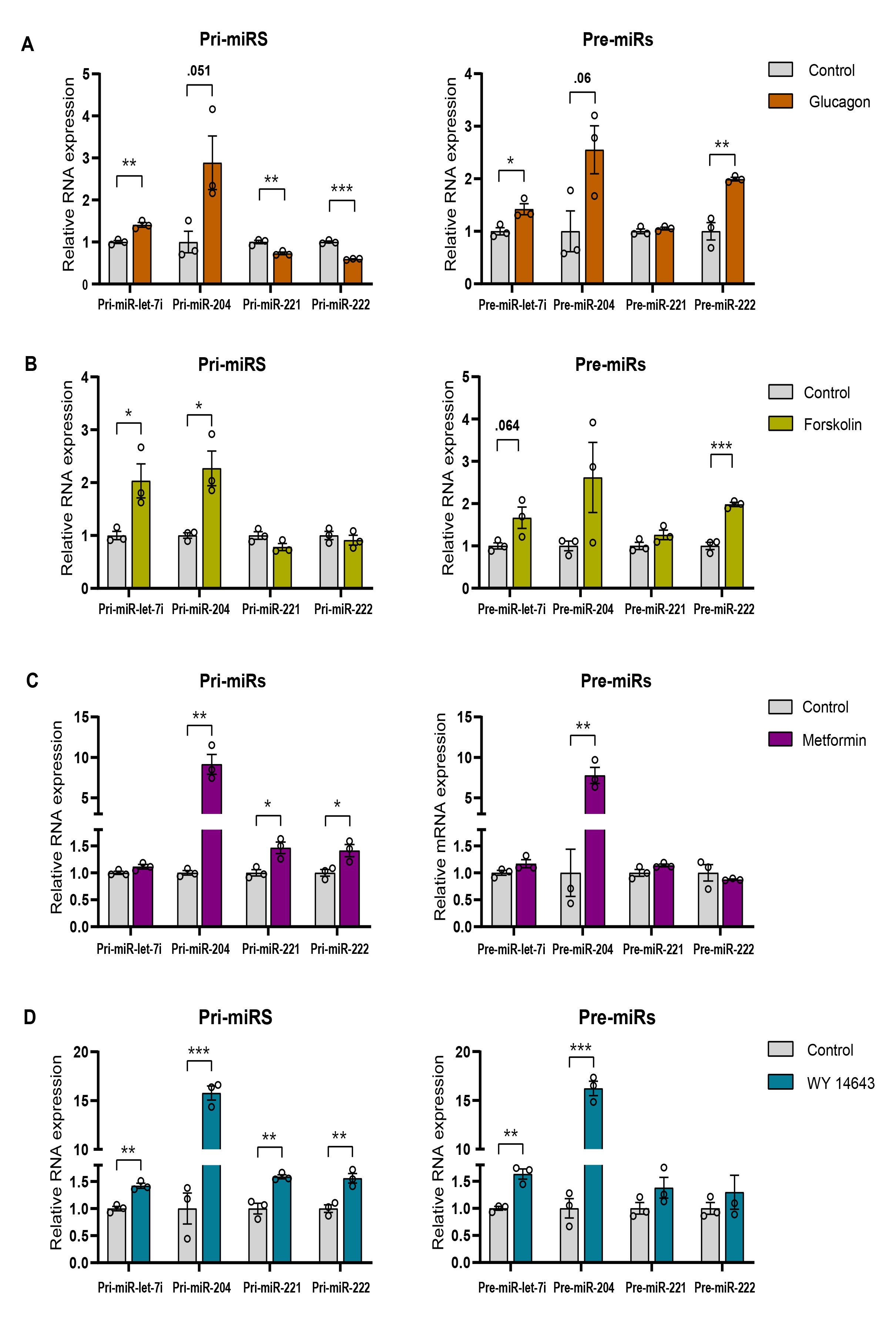
Starvation cues regulate miR-biogenesis in primary hepatocytes. **(A)** Relative expression of pri-miRs and pre-miR in primary hepatocytes treated with 30 nM of glucagon in LG media for 6 hrs. Pri-and pre-miR transcript levels were normalized to *Actin* mRNA level and plotted as fold change with respect to control (N=2, n=3). **(B)** Relative expression of pri-miRs and pre-miR in primary hepatocytes treated with 10 µM of forskolin in LG media for 12 hr. Pri-and pre-miR transcript levels were normalized to *Actin* mRNA level and plotted as fold change with respect to control (N=2, n=3). **(C)** Relative expression of pri-miRs and pre-miR in primary hepatocytes treated with 2 mM of metformin in HG media for 12 hrs. Pri-and pre-miR transcript levels were normalized to *Actin* mRNA level and plotted as fold change with respect to control (N=2, n=3). **(D)** Relative expression of pri-miRs and pre-miR in primary hepatocytes treated with 50 µM of WY-14643 in HG media for 24 hrs. Pri-and pre-miR transcript levels were normalized to *Actin* mRNA level and plotted as fold change with respect to control (N=2, n=3). Data information: LG-low glucose, HG-high glucose, data are represented as mean ± SEM. Statistical significance was calculated using a student t-test (*, p < 0.05; **, p < 0.01; ***, p < 0.001).

Next, we wanted to check if starvation-sensitive transcription factors are involved in the miR-biogenic program that we have described so far. Among others, PPARα is key for mediating the transcription of several hepatic genes in response to fasting. Hence, we treated primary hepatocytes with WY-14643, a specific agonist of PPARα, and found that the pri-and pre-levels were upregulated at both 10h (Fig S4B) and 24h (Fig 5C). Moreover, administration of Bezafibrate, which is used in the clinics, also resulted increased expression, phenocopying the effects observed with WY-14643 (Fig S4C). Together these findings clearly indicate that anticipatory expression of hepatic fed-miRs is associated with activation of metabo-endocrine factors, akin to starvation, and importantly hint at the potential of utilizing pharmacological agents to restore miR-homeostasis.

## DISCUSSION

Efficient physiological toggling between anabolic and catabolic pathways during fed-fast cycles is essential for metabolic homeostasis and an inability to do so has been associated with aging and non-communicable diseases (Ribarič, 2012; Geisler *et al*., 2016; Petersen, Vatner and Shulman, 2017; Paoli *et al*., 2019; -S. Designed Research; T and -S. Performed Research; T, 2023). Given that refeeding following starvation is often unanticipated, it is intuitive to expect that robust mechanisms would allow organisms to rapidly transit from fasted to a fed state (Jitrapakdee, 2012; Bideyan, Nagari and Tontonoz, 2021). While much is known about cephalic mechanisms (Power and Schulkin, no date b), molecular factors that govern gene expression programs and create anticipatory regulatory loops remain largely unknown. Further, owing to the fact that feeding and fasting are intricately coupled with light-dark or circadian cycles (Gerhart-Hines and Lazar, 2015; Reinke and Asher, 2019; Pickel and Sung, 2020b), it is important to delineate the roles of these zeitgebers in dictating molecular anticipation. In this context, we have unravelled the significance of metabolic and endocrine inputs in governing the anticipatory biogenesis of hepatic fed microRNAs. Importantly, we demonstrate the differential contribution of fed-fast and circadian cycles in controlling miR homeostasis in hepatocytes.

Post-transcriptional control of gene expression is known to provide energy-/time-efficient and dynamic spatio-temporal tuning of cellular functions (Halbeisen *et al*., 2008; Hocine, Singer and Grünwald, 2010). This is particularly relevant for fed-fast cycles since rapid and reversible inhibition of translation is essential (Koike *et al*., no date; Le Martelot *et al*., 2012). Consistent with this, others and we have demonstrated the importance of microRNAs in governing hepatic physiology (Hu *et al*., 2012; Du *et al*., 2014; Maniyadath *et al*., 2019). Strikingly, we also found that oscillatory fed-microRNAs, which inhibit the expression of catabolic genes and consequently enable fasted-refed transition, are expressed in anticipation during a starvation state (Maniyadath *et al*., 2019). Albeit parallel studies have also illustrated circadian dependence or rhythmicity in the expression of microRNAs across tissues, including the liver (Menet *et al*., 2012; Vollmers *et al*., 2012; Ji *et al*., 2023), whether fed-fast cycles contribute to miR biogenesis independently or cooperatively (with circadian inputs) remains unknown. In this context, we have delineated the causal upstream cues that dictate the anticipatory expression of hepatic fed-microRNAs, especially for let-7i, mir-204, mir-221, and mir-222, which not only displayed the most robust oscillation but also otherwise have been shown to be important for liver functions and hepatic cancers (Chen *et al*., 2014; Luo *et al*., 2017; Song *et al*., 2017b). Employing classical paradigms to perturb circadian and fed-fast cycles, we establish that while let-7i and mir-204 are tuned solely by fed-fast cues, mir-221 and mir-222 show only partial dependence. Specifically, comparing pri-miR levels in S-LD and S-DD showed that prolonged starvation induced heightened expression of let-7i and miR-204, to similar extents, clearly indicating that they are not dependent on circadian inputs. This also corroborated starvation dependence from AL-LD and S-LD paradigms.

miR processing has been shown to be regulated and several reports have indicated differential Dorsha/DGCR8 and Dicer activities to affect pri-, pre-and mature-miR levels (Ghatak and Sen, 2015; Creugny, Fender and Pfeffer, 2018; Vergani-Junior *et al*., 2021). Even though post-translational modifications, co-factors, RNA structures, and relative abundances of miR-transcripts have been proposed to impinge on Dorsha/DGCR8 and Dicer dependent maturation of pri-miRs to pre-and mature-miRs, respectively (Heo *et al*., 2008; Davis and Hata, 2009; Feng *et al*., 2011; Conrad *et al*., 2014; Alarcón *et al*., 2015; Louloupi *et al*., 2017), if/how this leads to global versus miR-specific effects remains less understood. This is relevant since we observed that while pre-miR-221/222 exhibited similar expression profile in *ad libitum* condition, they showed mutually exclusive pattern in response to continuous starvation. This differential response was conspicuous across all starvation paradigms and although intriguing, did suggest an intricate regulatory mechanism that determines miR-homeostasis, whose mechanistic basis will have to be unravelled in the future. An equally intriguing finding was fasted-refed transition dependent processing of pre-miRs to mature miRs, which is possibly mediated by anabolic inputs as an additional layer of regulation. Consistent with this, our results show that insulin signalling impinges on the maturation or processing of pri-/pre-miRs. Given that ERK, which is downstream of insulin signalling, has been otherwise shown to affect maturation of let-7i miRNA (Sun *et al*., 2016), we posit that upstream metabolic and neuroendocrine inputs associated with a refed state are required for hepatic miR maturation.

Another key highlight of the study is the observation of deregulated expression of hepatic fed miRs in response to dietary perturbations and aging. While qualitative and quantitative changes in miRs, intracellular and circulating, during aging and pathological conditions are well documented(‘Age-related changes in microRNA levels in serum’, no date; Jung and Suh, 2014; Deiuliis, 2016), whether these are resultant of altered miR biogenesis needs to be unravelled. In this context, we have observed that the biogenesis of microRNAs profiled in this study is significantly dampened in aging and over-nutrition paradigms. Notably, feeding mice with high-fat diet and sucrose water (along with a standard chow diet) muted the biogenesis of let-7i, miR-204, miR-221, and miR-222. Together these clearly suggest that both short-term and long-term dietary/metabolic inputs play a significant role in orchestrating miR homeostasis in the liver. It will be interesting to investigate, in the future, if such perturbations affect miR-biogenesis, degradation, and exosome-mediated secretion, globally.

Given the complex interplay between metabolism, miR-homeostasis, and organismal physiology, which are in turn dependent upon fed-fast and circadian cycles (Rottiers and Näär, 2012; Dumortier, Hinault and Van Obberghen, 2013; Hartig *et al*., 2015), it is nearly impossible to establish whether miR-biogenesis acts as an initiator or mediator mechanism. Nonetheless, dissecting molecular factors and evaluating the modulatory impact of commonly employed therapeutic interventions becomes crucial. In this regard, we not only provide mechanistic insights into key factors that link metabolic inputs to miR-biogenesis but also illustrate that, pharmacological agents, which regulate nodal molecular factors, impinge on hepatic fed-miR homeostasis. We clearly illustrate PPARα as one of the starvation dependent factors that exert control over the anticipatory expression of miRs. Importantly, treating hepatocytes with metformin and bezafibrate, which are well established to activate AMPK and PPARα dependent metabolic rewiring (Zhou *et al*., 2001; Puligheddu *et al*., 2013), led to a robust response vis-à-vis miR-biogenesis. Furthermore, taken together with our earlier report, the findings presented here raise the possibility of employing FDA-approved drugs to modulate miR expression to help mitigate physiological deficits associated with metabolic diseases.

In conclusion, our study unequivocally delineates fed-fast and circadian dependent expression of hepatic fed-miRs, which is pivotal for molecular anticipation. Employing paradigms that recapitulate normal physiological settings, we show complex interplay between metabolism and miR-biogenesis in a miR-dependent manner. Owing to the detrimental impact of modern lifestyles, largely mediated by perturbed fed-fast and circadian cycles (Arble *et al*., 2010; Charlot *et al*., 2021), we have unravelled physiological inputs that choreograph oscillatory/anticipatory microRNA biogenesis in the liver. Even though miRs have emerged as clinically feasible therapeutic targets, current approaches are based on miR-mimics and anti-miRs that affect individual microRNAs (Baumann and Winkler, 2014; Chakraborty *et al*., 2021; Diener, Keller and Meese, 2022). While this has merits, miRs are known to modulate cellular/organismal functions by exerting additive and convergent control over the expression of target mRNAs. Therefore, identifying physiological contexts and/or pharmacological interventions that reset miR-homeostasis, possibly by regulating a network of miRs as demonstrated in this study, will likely yield higher dividends.

## ACKNOWLEDGEMENTS

This research has been supported by the following funding sources: TIFR/DAE (19P0116) and Department of Biotechnology (BT/PR29878/PFN/20/1431/2018) to U.K.-S. We thank ACTREC Mumbai for providing us with C57BL/6NCrl mice. We thank the Advanced Research Unit on Metabolism, Development and Aging (ARUMDA) Consortium for supporting a part of this study. We especially thank Dr. Kalidas Kohale, Dr. Shital Suryavanshi, Chetan Sable, TIFR Mumbai and IISER Pune animal house staff for their help with the animal experiments. We extend our acknowledgement to UK lab members for their critical inputs and discussions during the study.

## AUTHOR CONTRIBUTIONS

Conceptualization, U.S.S. and U.K.-S.; methodology, U.S.S, S.C. and U.K.-S.; formal analysis, U.S.S, S.C. and U.K.-S.; investigation, U.S.S. and S.C.; writing, U.S.S., S.C., and U.K.-S.; funding acquisition, U.K.-S.

## DECLARATION OF INTERESTS

The authors declare no competing interest.

KEY RESOURCES TABLE

KEY RESOURCES TABLE
|  | SOURCE | IDENTIFIER |
| --- | --- | --- |
| **Chemicals, peptides and recombinant proteins** |  |  |
| RNaseOUT™ Recombinant Ribonuclease Inhibitor | Invitrogen | 10777019 |
| Random Hexamer (50uM) | Invitrogen | N8080127 |
| TriZol | Invitrogen | 15596018 |
| DEPC treated water | Invitrogen | AM9915G |
| UltraPure Distilled Water | Invitrogen | 10977015 |
| dNTP Mix | Genei | 652400021730 |
| Glucose Solution | Gibco™ | A2494001 |
| L- Glutamine | Sigma-Aldrich | 59202C |
| Sodium Pyruvate | Sigma-Aldrich | S8636 |
| Collagenase | Sigma-Aldrich | C5138 |
| Rat tail collagen | Sigma-Aldrich | C3867 |
| High Glucose DMEM | Sigma-Aldrich | D7777 |
| Low Glucose DMEM | Sigma-Aldrich | D5523 |
| DMEM without glucose | Sigma-Aldrich | D5030 |
| Fetal Bovine Serum | Gibco™ | 10270106 |
| Antibiotic Antimycotic Solution | Sigma-Aldrich | A5955 |
| Sodium pyruvate | Sigma-Aldrich | P225 |
| EGTA | Sigma-Aldrich | E3889 |
| Bovine serum albumin | MP Biomedicals | 160069 |
| DTT | Sigma-Aldrich | 578517 |
| Glucagon | Sigma-Aldrich | G2044 |
| HEPES | USB | 16926 |
| Forskolin | Sigma-Aldrich | F3917 |
| Metformin HCl | Cipla<br>Pharmaceuticals | - |
| WY-14643 | Sigma-Aldrich | C7081 |
| **Critical Commercial Assays** |  |  |
| SuperScript-IV RT kit | Invitrogen | 18090200 |
| KAPA SYBR® FAST Universal 2X qPCR<br>Master Mix | Roche | SFUKB |
| QuantimiR kit | System<br>Biosciences | Cat# RA420A-1 |
| **Experimental Models: Organisms/Strains** |  |  |
| Mouse: C57BL/6NCrI | CRL | Strain No: 027 |

