## Supplementary figures and images for "Anticipatory Biogenesis of Hepatic Fed MicroRNAs is Regulated by Metabolic and Circadian Inputs"

### Supplementary Figures

##### SUPPLEMENTARY FIGURE 1

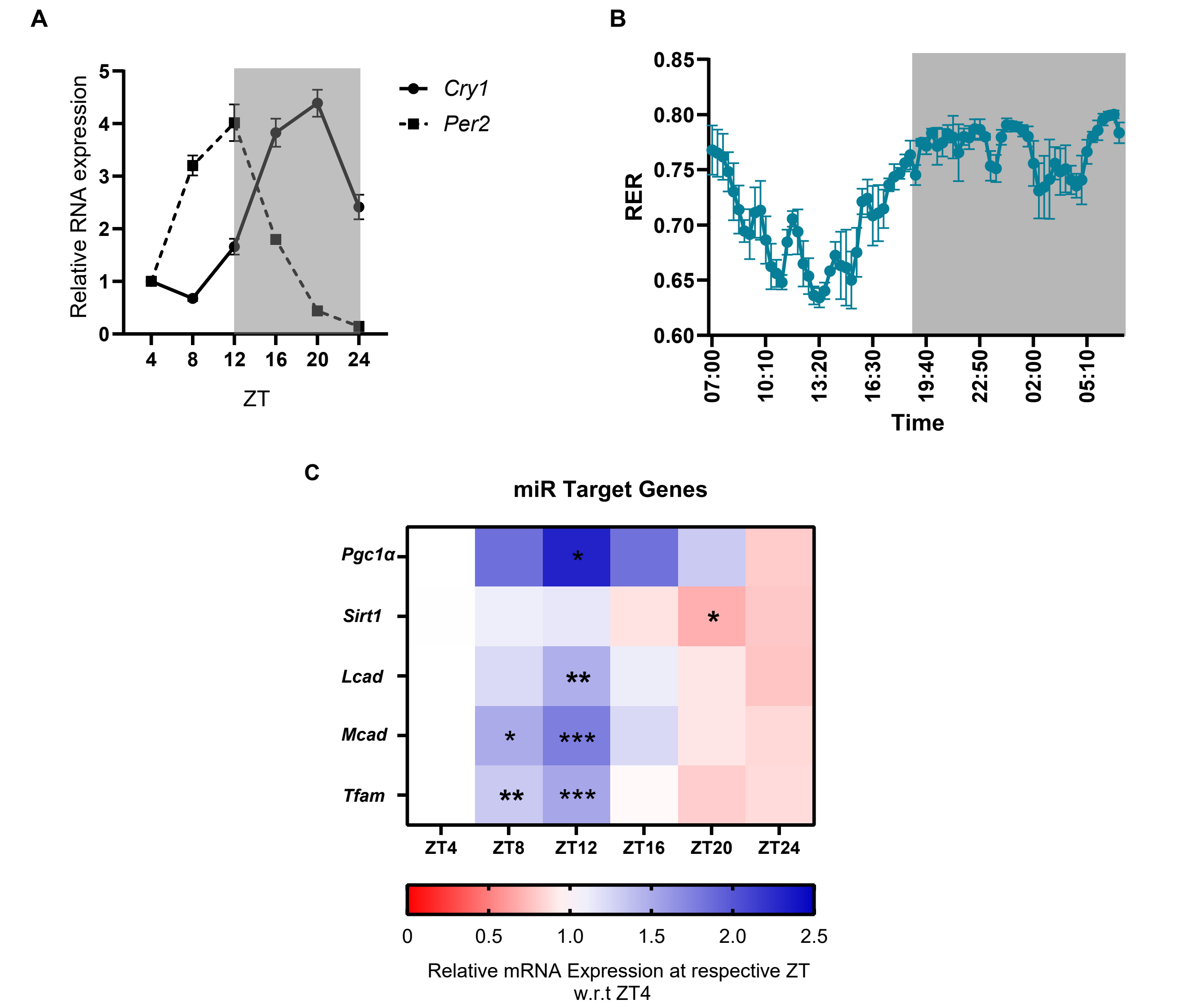

#### SUPPLEMENTARY FIGURE 2

A

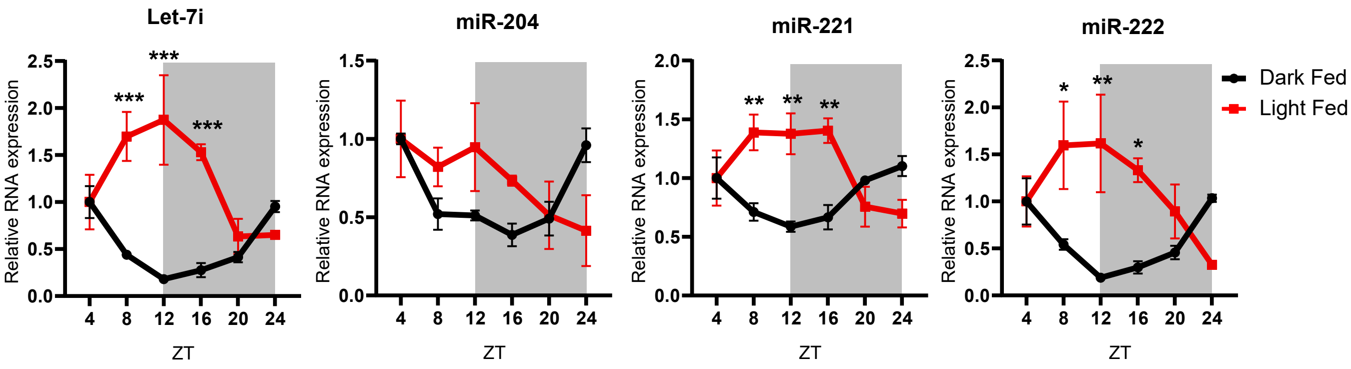


B

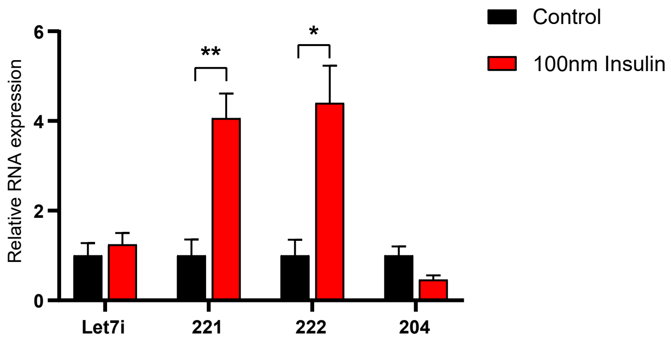

#### SUPPLEMENTARY FIGURE 3

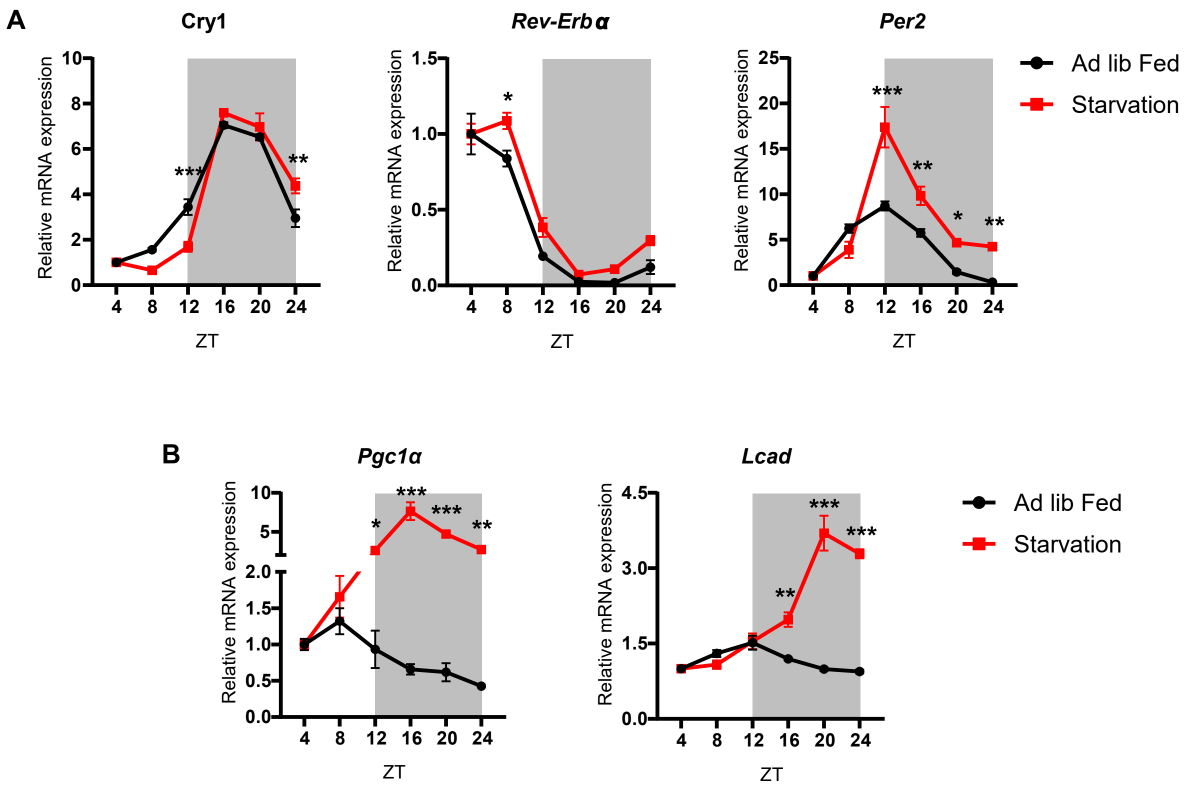

##### SUPPLEMENTARY FIGURE 4

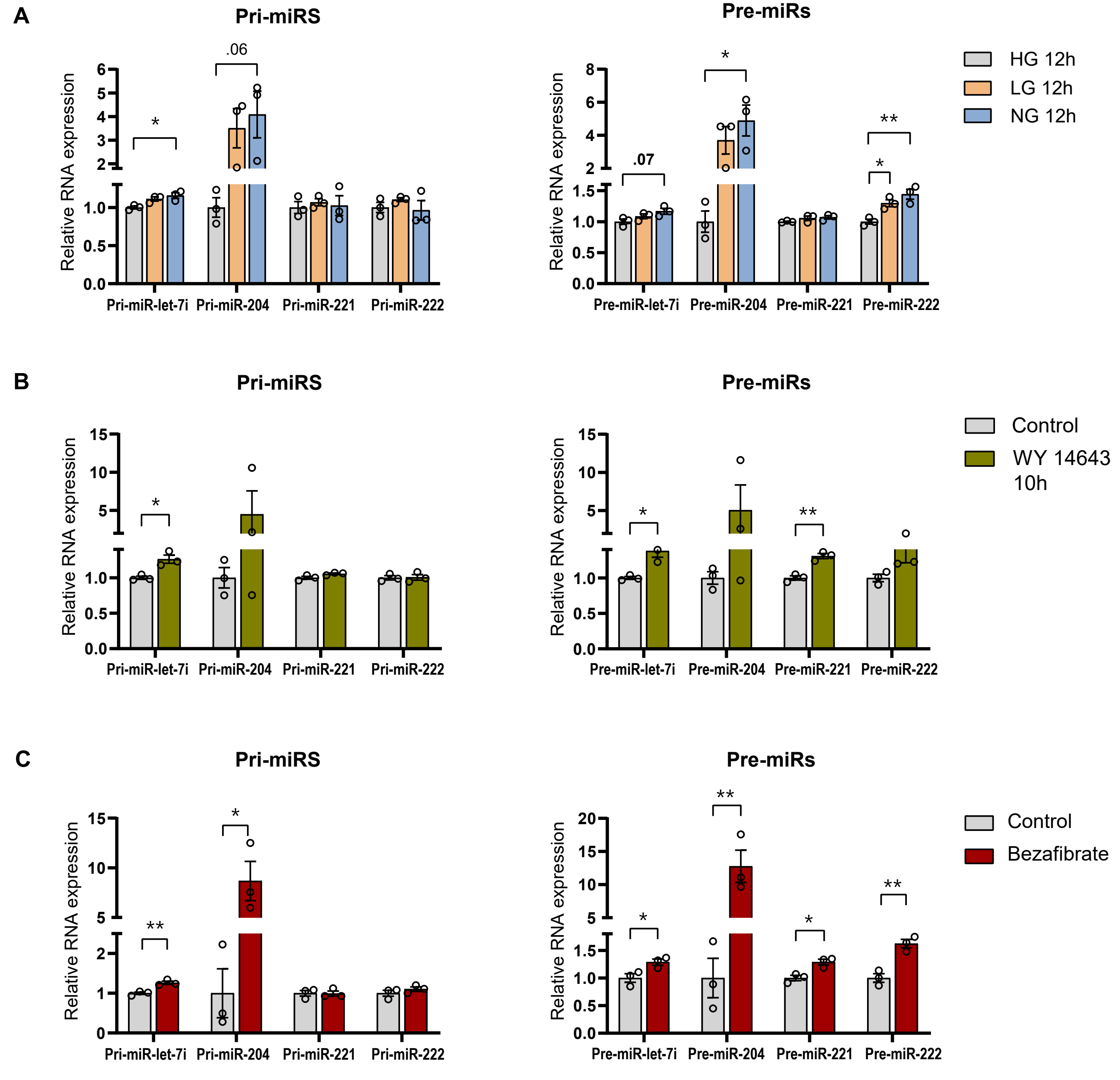
