## Supplemental Information for "Anticipatory Biogenesis of Hepatic Fed MicroRNAs is Regulated by Metabolic and Circadian Inputs"

**Supplementary Figure Legends**

**Figure S1: Diurnal rhythmicity of miRNA targets (starvation-dependent mRNAs) in liver**

**(A**) Relative expression of *Cry1* and *Per2* from liver of *ad libitum* fed mice at the indicated ZT. *Cry1* and *Per2* mRNA levels were normalized to *Actin* mRNA level and plotted as fold change with respect to ZT4 (N=2, n=3).

**(B)** Respiratory exchange ratio of *ad libitum* fed mice over a 24-hr light-dark cycle (n=3)

**(C)** Heatmap depicting relative expression of the indicated target starvation-responsive mRNA transcripts at the respective ZT. mRNA levels of the respective genes were normalized to *Actin* mRNA level and plotted as fold change with respect to ZT4 (N=2, n=3).

Data are represented as mean ± SEM. Statistical significance was calculated using one-way ANOVA with Tukey's test for multiple comparisons between groups (*, $p < 0.05$; **, $p < 0.01$; ***, $p < 0.001$).

**Figure S2: Temporal feeding paradigms reprograms oscillatory hepatic fed-miRNAs and is dependent on insulin**

**(A)** Relative expression of mature miRs from the liver of dark-fed and light-fed fed mice at the indicated ZT. Mature miR transcript levels were normalized to 18S rRNA level and plotted as fold change with respect to ZT4 (N=2, n=3).

**(B)** Relative expression of mature miRs in primary hepatocytes treated with 100 nM insulin for 1.5 hrs. Mature miR transcript levels were normalized to 18S rRNA level and plotted as fold change with respect to control. (N=2, n=3).

Data are represented as mean ± SEM. For **A**, statistical significance between dark-fed and light-fed group at indicated ZT was calculated using multiple t-tests with Holm-Sidak correction. For **B**, statistical significance was calculated using students t-test. (*, $p < 0.05$; **, $p < 0.01$; ***, $p < 0.001$).

**Figure S3: Hepatic clock components sustain oscillations in starvation but not fasting genes**

**(A)** Relative expression of liver clock genes *Cry1*, *Rev-Erbα*, and *Per2* from the liver of AL-DD and S-DD mice, at the indicated ZT. Respective gene mRNA levels were normalized to *Actin* mRNA level and plotted as fold change with respect to ZT4 (N=2, n=3).

**(B)** Relative expression of liver clock genes *Pgc1α*, and *Lcad* from the liver of AL-DD and S-DD mice, at the indicated ZT. Respective gene mRNA levels were normalized to *Actin* mRNA level and plotted as fold change with respect to ZT4 (N=2, n=3).

Data information: AL-LD - *ad libitum* light-dark, S-LD - starvation light-dark, data are represented as mean ± SEM. Statistical significance between *ad libitum* fed and progressively starved mice at indicated ZT was calculated using multiple t-tests with Holm-Sidak correction (*, $p < 0.05$; **, $p < 0.01$; ***, $p < 0.001$).

**Figure S4: Starvation cues regulate miR-biogenesis in primary hepatocytes**

**(A)** Relative expression of pri-miRs and pre-miR in primary hepatocytes treated with HG, LG, NG media for 12 hrs. Pri- and pre- miR transcript levels were normalized to *Actin* mRNA level and plotted as fold change with respect to control (N=2, n=3).

**(B)** Relative expression of pri-miRs and pre-miR in primary hepatocytes treated with 100 µM of bezafibrate in LG media for 14 hrs. Pri- and pre- miR transcript levels were normalized to *Actin* mRNA level and plotted as fold change with respect to control (N=2, n=3).

Data information: HG- high glucose, LG- low glucose, NG- no glucose, data are represented as mean ± SEM. Statistical significance was calculated using a student t-test (*, $p < 0.05$; **, $p < 0.01$; ***, $p < 0.001$)

***Table S1: Primer sequences for RT-qPCR -***

| GENES | PRIMER SEQUENCE |
|---|---|
| *Sirt1* | FP:5'-GTAACCCTGTAAAGCTTTCAG-3'<br>RP:5'-CAGAAGAGTCTTGTGGTACAG-3' |

| | |
|---|---|
| *Mcad* | FP:5'-TTGAGTTCACCGAACAGCAG-3'<br>RP:5'-ATCCGCTGCACAGATCCAAA-3' |
| *Lcad* | FP:5'-TCTTTTCCTCGGAGCATGACA-3'<br>RP:5'-GACCTCTCTACTCACTTCTGA-3' |
| *Ppargc1a* | FP:5'-GTGGATGAAGACGGATTGCC-3'<br>RP:5'-GCTGAGTGTTGGCTGGTGCC-3' |
| *Tfam* | FP:5'-CAAGTCAGCTGATGGGTATGG-3'<br>RP:5'-TTTCCCTGAGCCGAATCATCC-3' |
| *Cry1* | FP:5'-CACTGGTTCCGAAAGGGACTC-3'<br>RP:5'-CTGAAGCAAAAATCGCCACCT-3' |
| *Per2* | FP:5'-CTCCAGCGGAAACGAGAACT-3'<br>RP:5'-CTCACTACTGCAGCCGCTC-3' |
| *Rev-Erbα* | FP:5'-TGGCATGGTGCTACTGTGTAAGG-3'<br>RP:5'-ATATTCTGTTGGATGCTCCGGCG-3' |
| *Pri-miR-Let7i* | FP:5'-TCCGCCGGCTCCCACACCAT-3'<br>RP:5'-CGCGGCGCTGAGCATCACCA-3' |
| *Pri-miR-204* | FP:5'-TCTTCATGTGACTCGTGGAC-3'<br>RP:5'-AGATGCCAGTGATGACAATTG-3' |
| *Pri-miR-221* | FP:5'-CTGTATGGATAATTTGCAGGC-3'<br>RP:5'-GGCCAATGAACAAATGATTCC-3' |
| *Pri-miR-222* | FP:5'-ACTCTCACAAAGGATTAGGGTG-3'<br>RP:5'-GGGTACCATAACTCAAGCTAG-3' |
| *Pre-miR-Let7i* | FP:5'-CTGGCTGAGGTAGTAGTTTG-3'<br>RP:5'-TAGCAAGGCAGTAGCTTGCG-3' |

| | |
|---|---|
| *Pre-miR-204* | FP:5'-GGCTACAGTCTTTCTTCATGT-3'<br>RP:5'-GCCAGTGATGACAATTGAACG-3' |
| *Pre-miR-221* | FP:5'-GAGAACATGTTTCCAGGTAGC-3'<br>RP:5'-TGAACATCCAGATCTGGG-3' |
| *Pre-miR-222* | FP:5'-GCTGCTGGAAGGTGTAGG-3'<br>RP:5'-GCTAGAAGATGCCATCAGAG-3' |
| *Actb* | FP:5'-CCTCCCTGGAGAAGAGCTATGA-3'<br>RP:5'-GCACTGTGTTGGCATAGAGGTC-3' |
| 18S | FP:5'-TTTCGAGGCCCTGTAATTGG-3' |
| Mature Let7i | FP:5'-TGAGGTAGTAGTTTGTGCTGTT-3' |
| Mature miR-221 | FP:5'-AGCTACATTGTGGCTGGGTTTC-3' |
| Mature miR-222 | FP:5'-AGCTACATCTAGCTACTGGGT-3' |
| Mature miR-204 | FP:5'-TTCCCTTTGTCATCCTATGCCT-3' |
